# Developmental Dynamics of Phenotypic Architecture in ASD

**DOI:** 10.64898/2026.05.25.727621

**Authors:** Alejandra Fernandez

## Abstract

The behavioral features of autism spectrum disorders (ASD) span multiple domains, yet how behavioral dimensions shape and contribute to ASD phenotypes remains unknown. Here, using large-scale phenotypic data from SPARK, I identified a reproducible architecture linking sensory-repetitive, motor, and social-communication features with broader clinical variation and show broad associations with psychopathology. This organization is recovered across independent SPARK subsets and independently replicated in the Simons Simplex Collection, where sensory-repetitive features again form the highest-ranking hubs. Anxiety-related symptoms contribute to but not fully account for this organization. Age-stratified analyses in SPARK revealed a marked reorganization of phenotypic architecture with sensory-repetitive features dominating the hub hierarchy in childhood and motor features rising to the highest relative positions in adolescence and adulthood. Together, these findings suggest that autism-related heterogeneity is not simply multidimensional but organized around a reproducible sensory-repetitive-centered architecture whose developmental expression varies across cohorts.

**Highlights:**

- Features that define autism are not necessarily those that organize its phenotype
- Sensory-repetitive features form a reproducible center of phenotypic architecture
- Anxiety shapes but does not account for sensory-repetitive centrality
- Overall architecture replicates across cohorts

**Graphical abstract:** 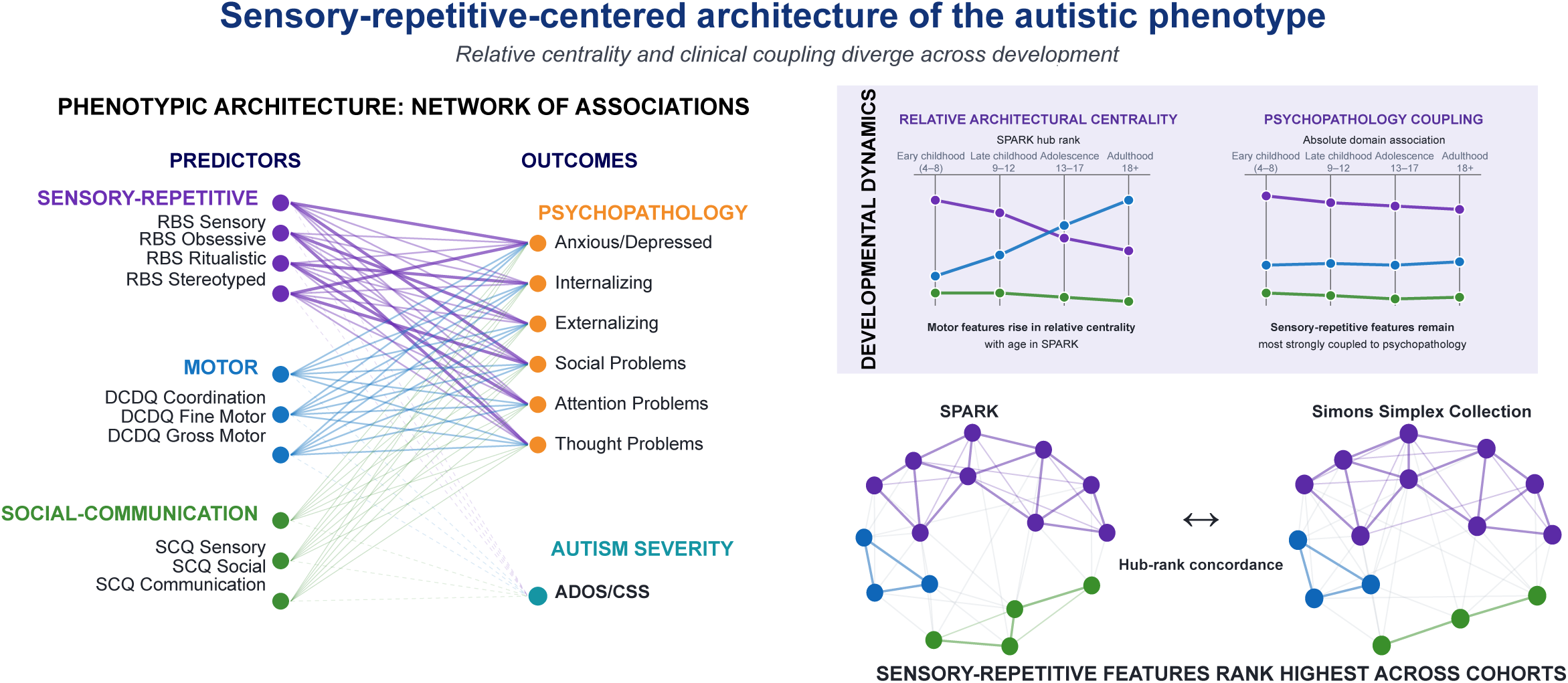

## Introduction

Autism spectrum disorder (ASD) is defined by differences in social communication and restricted or repetitive behaviors, yet its clinical phenotype extends far beyond the features that establish diagnosis, and include substantial behavioral heterogeneity spanning motor, sensory, repetitive, social-communicative, and cognitive domains [1–3]. While typically measured and studied in isolation, these features do not occur independently. Motor difficulties co-occur with social and behavioral differences, sensory features are associated with repetitive behavior and psychopathology, and behavioral difficulties often extend across multiple clinical and symptom domains [4–9]. Whether ASD-associated behavioral features simply co-occur or form a structured architecture remains an open question, and raises the possibility of a distinction between diagnostic prominence and phenotypic centrality. Neither diagnostic criteria nor severity measures quantify how deeply embedded features are within the broader behavioral phenotype, as they aim to define a disorder and measure prevalence of particular features respectively [10]. A feature could be diagnostically salient yet relatively circumscribed, whereas another display architectural centrality: the extent to which a behavioral feature shows broad and strong associations across the phenotype, shifting the focus from most characteristic features to the most connected to the broader ASD clinical phenotype [11–13].

Sensory and repetitive behaviors are particularly compelling candidates for such a role. Sensory differences emerge early in development and can persist throughout childhood [14, 15], with early sensory hyperreactivity associated with later emotional difficulties [16]. Sensory and repetitive features are also closely linked to anxiety and internalizing symptoms, although these relationships appear to involve partially separable pathways [17, 18]. Moreover, repetitive sensory-motor behaviors have further been associated with subsequent social development [19]. while these observations are suggestive of a cross-domain connection in the phenotype, rather than isolated comorbidities the position of sensory-repetitive features within the broader phenotypic architecture has not been established. How central these features are remains unclear, and how phenotypic organization evolves remains to be determined. As behavioral architecture is not static throughout development [20–23], distinguishing developmental shifts may help explain why different behavioral features become clinically prominent at different stages while other cross-domain relationships remain persistent.

Here, I used the large SPARK cohort [24] to map cross-domain phenotypic architecture in ASD to identify behavioral features occupying central positions within the phenotype. I found that this architecture was reproducible within SPARK and generalizes to the independent Simons Simplex Collection (SSC) [25]. Moreover, this architecture persists after removing anxiety-related variance and seems to evolve throughout development. A consistent organizing principle emerged, highlighting centrality of sensory-repetitive features in the autistic behavioral phenotype and showing particularly strong relationships with psychopathology. Developmental analyses further revealed that relative hub organization and absolute psychopathology coupling are not interchangeable: motor features became more prominent within the SPARK hierarchy with age, whereas sensory-repetitive features remained the domain most strongly coupled to psychopathology.

Together, these findings shed light into the organized and developmentally structured phenotypic architecture in ASD.

## Results

### Cross-Domain Associations Within Autism Phenotypic Architecture

I first asked whether behavioral features spanning motor coordination, sensory-repetitive behavior (SR), and social-communication domains showed systematic cross-domain associations with psychopathology and clinician-rated autism severity. This analysis was intended to establish the overall organization of the phenotype before testing whether specific behavioral features functioned as disproportionately connected “hubs.” I examined 10 behavioral predictors derived from the Developmental Coordination Disorder Questionnaire (DCDQ), Repetitive Behavior Scale–Revised (RBS-R), and Social Communication Questionnaire (SCQ) against nine outcomes spanning parent-reported psychopathology, from six Child Behavior Checklist (CBCL) scales, and clinician-observed autism symptom severity, from three Autism Diagnostic Observation Schedule (ADOS) calibrated severity scores (CSS) (**Figure 1**). For each predictor-outcome combination, I quantified the predictor’s incremental contribution beyond age, sex, and nonverbal IQ using signed √ΔR². Specifically, I compared a covariate-only model with a model additionally containing the behavioral predictor, calculated the increase in explained variance attributable to the predictor, and assigned the sign of the predictor’s regression coefficient to the resulting √ΔR² [26–28]. Thus, the magnitude of signed √ΔR² reflects the predictor’s unique contribution beyond the covariates, while its sign indicates the direction of the adjusted association. Analyses used all available observations for each predictor-outcome combination, such that sample size varied across cells. Across the resulting 90 predictor-outcome tests, 60 survived FDR correction (q <.05).

**Figure 1.**
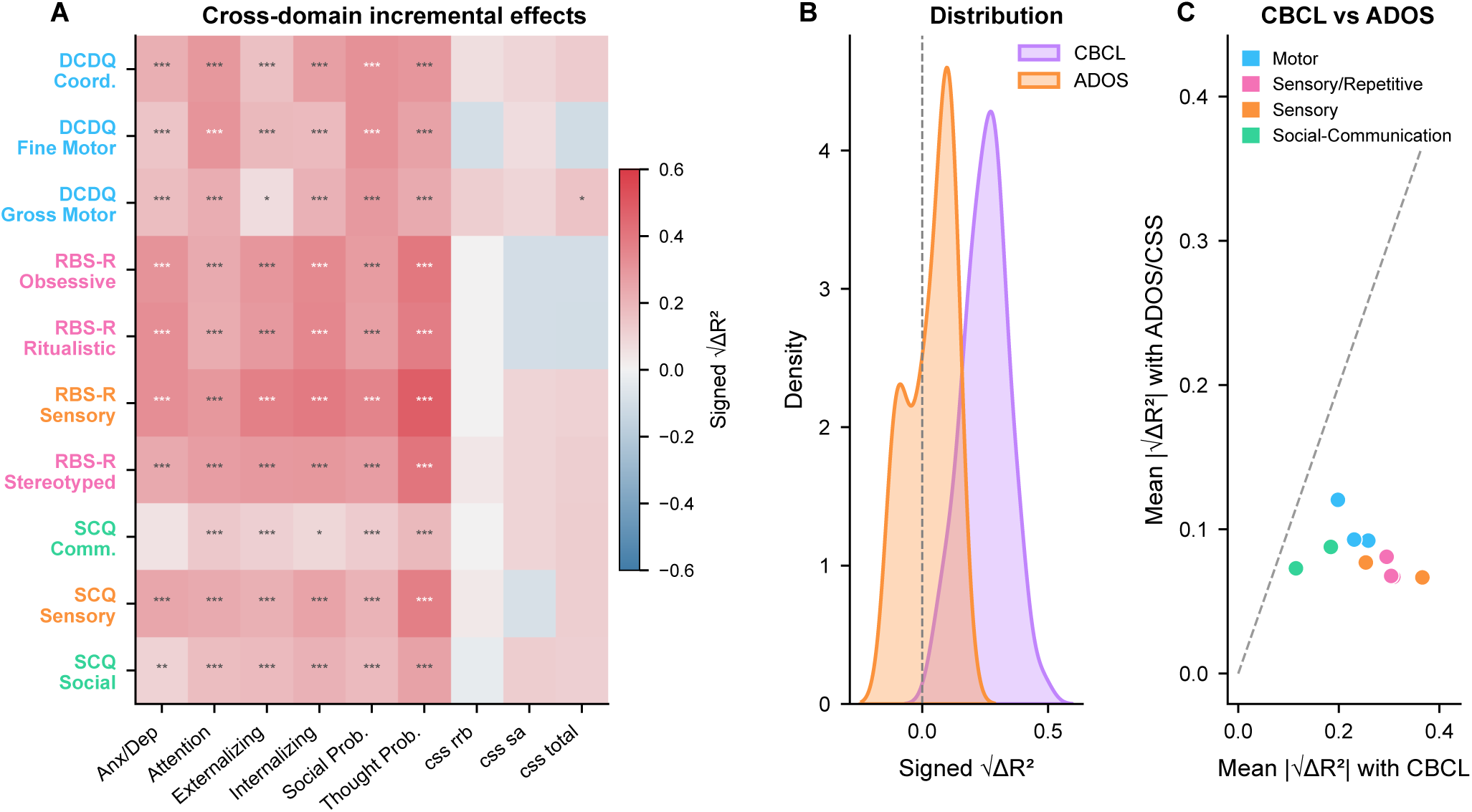
Cross-domain phenotypic architecture in autism spectrum disorder. Covariate-adjusted associations between 10 behavioral predictors spanning motor (DCDQ), sensory-repetitive (RBS-R), and social-communication (SCQ) domains and nine outcomes encompassing CBCL psychopathology and clinician-rated autism severity (ADOS CSS) in SPARK. Associations are expressed as signed √ΔR², defined as sign(β) × √max(0, R²_full − R²_base), where the baseline model included age, sex, and NVIQ. Benjamini–Hochberg FDR correction was applied across the 90 predictor × outcome tests. **(A)** Predictor × outcome association matrix. Color encodes signed √ΔR², with significance markers identifying associations surviving FDR correction. Associations were widespread across CBCL psychopathology outcomes but substantially weaker for ADOS CSS outcomes; 59 of 60 CBCL associations and 1 of 30 ADOS CSS associations survived FDR correction. **(B)** Distribution of signed √ΔR² estimates for associations with CBCL versus ADOS CSS outcomes, illustrating the substantially larger effects involving psychopathology. **(C)** Mean absolute √ΔR² for each behavioral predictor across CBCL outcomes versus ADOS CSS outcomes. The dashed identity line denotes equal average association magnitude with the two outcome domains.

The resulting matrix revealed extensive but nonuniform cross-domain coupling (**Figure 1A**). Incremental effects were particularly prominent among sensory-repetitive predictors and CBCL psychopathology outcomes. Among all predictor-outcome pairs, the strongest association was between RBS-R Sensory and CBCL Thought Problems (√ΔR² =.478, n = 1,064, q <.001). RBS-R Sensory also showed substantial incremental effects for CBCL Internalizing (√ΔR² =.387, n = 1,082, q <.001), Externalizing (√ΔR² =.366, n = 1,082, q <.001), and Social Problems (√ΔR² =.352, n = 1,064, q <.001). This pattern was not limited to the sensory subscale: RBS-R Stereotyped behavior showed a √ΔR² of.405 with CBCL Thought Problems, and RBS-R Obsessive behavior showed a √ΔR² of.398 with the same outcome (both q <.001). Motor predictors showed cross-domain effects of intermediate magnitude. DCDQ Coordination and DCDQ Fine Motor, for example, showed √ΔR² values of.314 and.311, respectively, with CBCL Social Problems (both n = 1,030, q <.001). In contrast, SCQ Communication generally showed smaller incremental effects across the matrix; among CBCL outcomes, its effects ranged from √ΔR² =.048 for Anxious/Depressed (q =.286) to.185 for Thought Problems (n = 1,062, q <.001). Thus, behavioral dimensions were not equivalently connected to the broader phenotype: some features contributed unique information across multiple outcomes, whereas others showed a substantially more restricted pattern of association. This heterogeneity provided the initial motivation for quantifying cross-domain hubness in the subsequent analysis.

The architecture also differed markedly by outcome domain (**Figure 1B**). √ΔR² values for CBCL outcomes were predominantly positive and shifted toward larger effect magnitudes, whereas effects involving ADOS/CSS were substantially smaller and included a negative tail. Negative √ΔR² values indicate inverse predictor-outcome associations after adjustment for age, sex, and nonverbal IQ, rather than negative explained variance. Across the 60 predictor-CBCL combinations, the mean √ΔR² was.251, compared with.034 across the 30 predictor-ADOS/CSS combinations. Fifty-nine of 60 CBCL associations survived FDR correction, whereas only one of 30 ADOS/CSS associations did. Thus, the difference between outcome domains was not driven by a small number of individual associations: behavioral predictors contributed substantially more unique explanatory information to variation in parent-reported psychopathology than to clinician-rated autism symptom severity.

I next asked whether this difference was broadly shared across predictors or driven by a small subset of behavioral features (**Figure 1C**). For each predictor, I calculated the mean absolute signed √ΔR² across the six CBCL outcomes and separately across the three ADOS/CSS outcomes. Taking the absolute value at this stage allowed this panel to compare the magnitude of incremental effects irrespective of direction. The identity line represents equal average effect magnitude in the two outcome domains; points below the line therefore indicate stronger average incremental effects for CBCL than for ADOS/CSS. All predictors fell below this line, demonstrating that the preferential association with psychopathology was distributed across behavioral domains rather than attributable to a single predictor. The contrast was especially pronounced among RBS-R predictors. RBS-R Sensory had a mean absolute √ΔR² of.366 across CBCL outcomes compared with.067 across ADOS/CSS outcomes, while RBS-R Obsessive and Ritualistic behaviors similarly showed large mean CBCL effects. Motor predictors generally occupied an intermediate position. DCDQ Coordination, for example, showed a mean absolute √ΔR² of.259 across CBCL outcomes compared with.092 across ADOS/CSS outcomes. Social-communication predictors generally showed smaller cross-domain effect magnitudes, with SCQ Communication displaying the smallest mean CBCL effect.

Together, these analyses establish that associations are widespread but strongly heterogeneous across behavioral predictors, suggesting that some dimensions may occupy more central positions in the phenotype than others. I therefore next quantified predictor-level hubness across outcomes to identify which behavioral dimensions most consistently contribute unique explanatory information across the broader phenotypic architecture.

### Phenotypic Architecture Is Organized Around Sensory-Repetitive Dimensions

Having established that cross-domain incremental effects were widespread but heterogeneous across behavioral dimensions, I next asked whether some predictors were consistently more strongly connected to the broader phenotype than others. I quantified this property using a Hubness Index that integrates both the breadth and magnitude of each predictor’s cross-domain associations. For each behavioral dimension, the index was calculated as the sum of the absolute √ΔR² values across outcomes surviving FDR correction (*q* <.05), after adjustment for age, sex, and nonverbal IQ. Thus, high hubness reflects both the number of significant cross-domain associations and the magnitude of these. Hubness revealed a pronounced hierarchy across the 10 behavioral dimensions (**Figure 2A**). RBS-R Sensory showed the highest overall connectivity (Hubness Index = 2.20), approximately 3.4-fold greater than SCQ Communication, which ranked lowest (0.64). Sensory-repetitive dimensions dominated the upper end of the hierarchy: RBS-R Sensory (2.20), Obsessive (1.85), Ritualistic (1.82), and Stereotyped (1.77) occupied ranks 1–4, with each showing significant associations with 6 of 9 outcomes. DCDQ Coordination (1.55, rank 5) and SCQ Sensory (1.52, rank 6) formed an intermediate tier, followed by DCDQ Fine Motor (1.38, rank 7), DCDQ Gross Motor (1.34, rank 8), and SCQ Social (1.10, rank 9). SCQ Communication was markedly less connected than the remaining dimensions (0.64, rank 10), with significant associations with 5 of 9 outcomes. The resulting ordering therefore did not simply separate instruments: although RBS-R dimensions predominated among the highest-ranked predictors, sensory, motor, and social-communication features were distributed across the remainder of the hierarchy.

**Figure 2.**
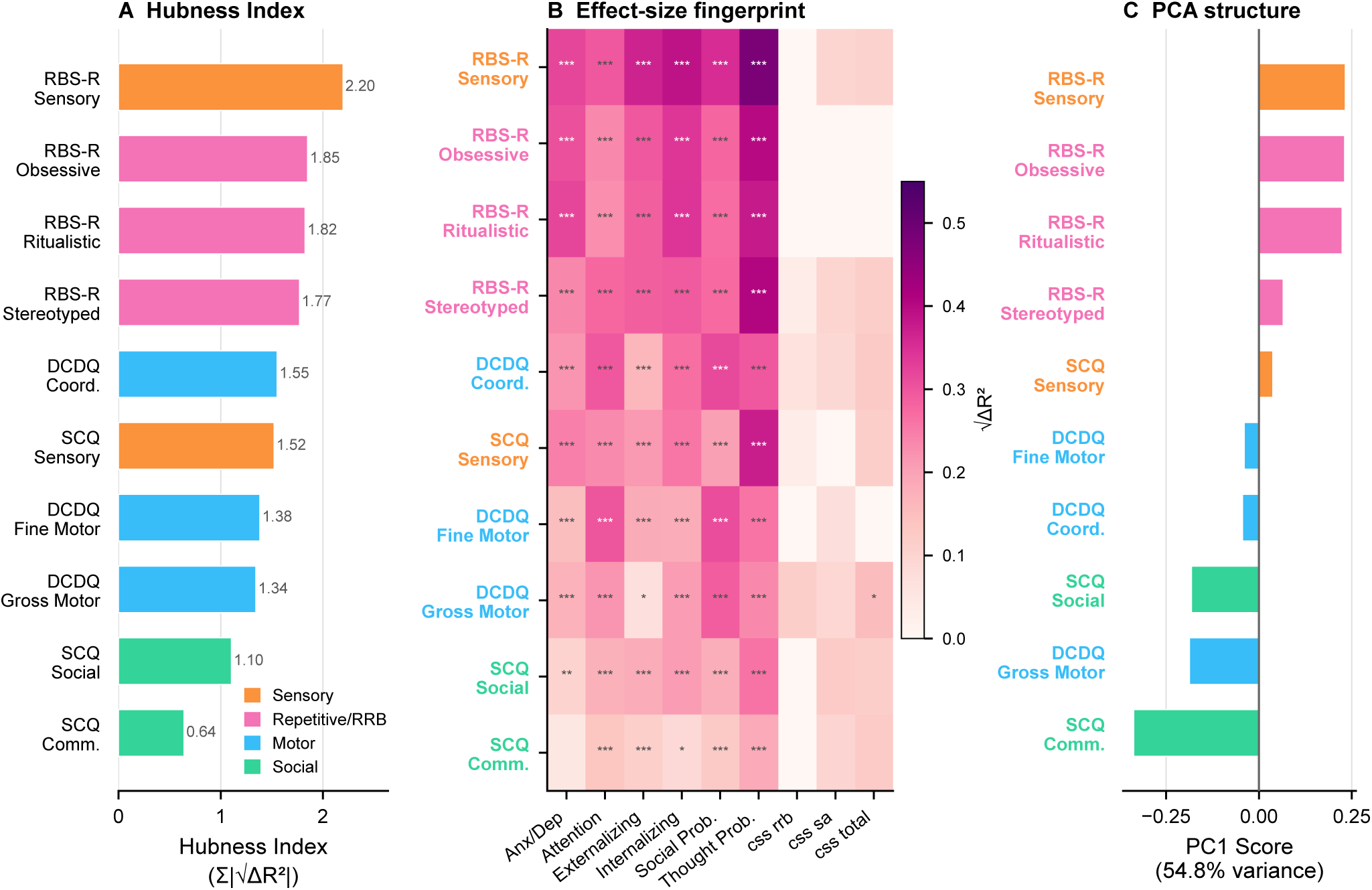
Sensory-repetitive dimensions function as phenotypic hubs. Hub organization was derived from the covariate-adjusted signed √ΔR² architecture shown in Figure 1. Predictor labels are color-coded by behavioral domain. **(A)** Hubness Index for each behavioral predictor, defined as the sum of |√ΔR²| across its FDR-significant outcome associations (*q* <.05). RBS-R Sensory showed the greatest hubness (H = 2.20), followed by RBS-R Obsessive (H = 1.85), Ritualistic (H = 1.82), and Stereotyped (H = 1.77). **(B)** Covariate-adjusted effect-size fingerprint underlying the hub hierarchy. Each cell represents signed √ΔR² for one predictor × outcome association; significance markers denote FDR-corrected associations. RBS-R Sensory × CBCL Thought Problems showed the largest individual effect (√ΔR² =.478). **(C)** Predictor scores on the first principal component of the association architecture. PC1 explained 54.8% of variance, with sensory-repetitive dimensions concentrated toward the positive end of the dominant architectural axis.

I next examined the predictor-outcome fingerprints underlying these aggregate hub scores (**Figure 2B**). The √ΔR² matrix showed that high hubness among RBS-R dimensions was driven primarily by broad effects across the six CBCL psychopathology outcomes rather than by a single exceptionally strong association. RBS-R Sensory displayed the strongest overall fingerprint, including the largest individual effect in the matrix for CBCL Thought Problems (√ΔR² =.478), and substantial effects across the remaining CBCL scales. RBS-R Obsessive and Stereotyped behaviors likewise showed large effects for Thought Problems (√ΔR² =.398 and.405, respectively), while Ritualistic behavior showed a similarly broad CBCL profile. Motor dimensions generally showed effects of intermediate magnitude, whereas SCQ Communication displayed the weakest overall fingerprint. The fingerprint matrix also clarified why overall hubness differed strongly across predictors.

Hubness was driven predominantly by the accumulation of effects across psychopathology outcomes: 59 of the 60 predictor-CBCL associations survived FDR correction, whereas only 1 of the 30 predictor-ADOS/CSS associations did. The sole FDR-significant ADOS/CSS effect was DCDQ Gross Motor × CSS Total (signed √ΔR² =.154, *n* = 375, *q* =.040). Thus, the high ranking of sensory-repetitive dimensions reflects broad covariate-adjusted connectivity with multiple forms of psychopathology rather than uniformly elevated associations across all outcome domains. This pattern is consistent with the CBCL–ADOS/CSS separation established in **Figure 1**.

Finally, I asked whether the same predictor hierarchy emerged from the multivariate structure of the association matrix without explicitly defining hubness. Principal component analysis of the predictor × outcome signed √ΔR² matrix identified a dominant first component accounting for 54.8% of variance in cross-domain association profiles (Figure 2C). RBS-R dimensions occupied the positive end of PC1, led by Sensory (PC1 = 3.13), Obsessive (2.30), Ritualistic (2.18), and Stereotyped (1.01). SCQ Communication occupied the opposite extreme (−4.38), followed by SCQ Social (−2.34) and DCDQ Gross Motor (−1.92), while DCDQ Coordination was positioned near the center of the axis. The second principal component accounted for an additional 19.6% of variance and captured variation in association profiles not represented by the dominant PC1 gradient. The convergence of these complementary representations indicates that sensory-repetitive centrality is not an artifact of a single summary metric. The Hubness Index identifies a strong rank ordering in cross-domain connectivity (**Figure 2A**); the underlying fingerprint matrix shows that this ordering arises primarily from the extent and magnitude of associations with psychopathology outcomes (**Figure 2B**); and an unsupervised decomposition of the association matrix independently identifies a dominant axis along which sensory-repetitive dimensions occupy one extreme and weakly connected social-communication dimensions the other (**Figure 2C**). Together, these findings identify sensory-repetitive features, particularly RBS-R Sensory, as central organizing dimensions of the cross-domain phenotypic architecture.

### Phenotypic Architecture Is Internally Reproducible

I next asked whether the phenotypic architecture identified in the full SPARK sample could be recovered independently from nonoverlapping subsets of participants.

Reproducibility was evaluated at multiple levels, from individual predictor-outcome effects to the higher-order hub hierarchy and domain-level architecture (**Figure 3**) [29].

**Figure 3.**
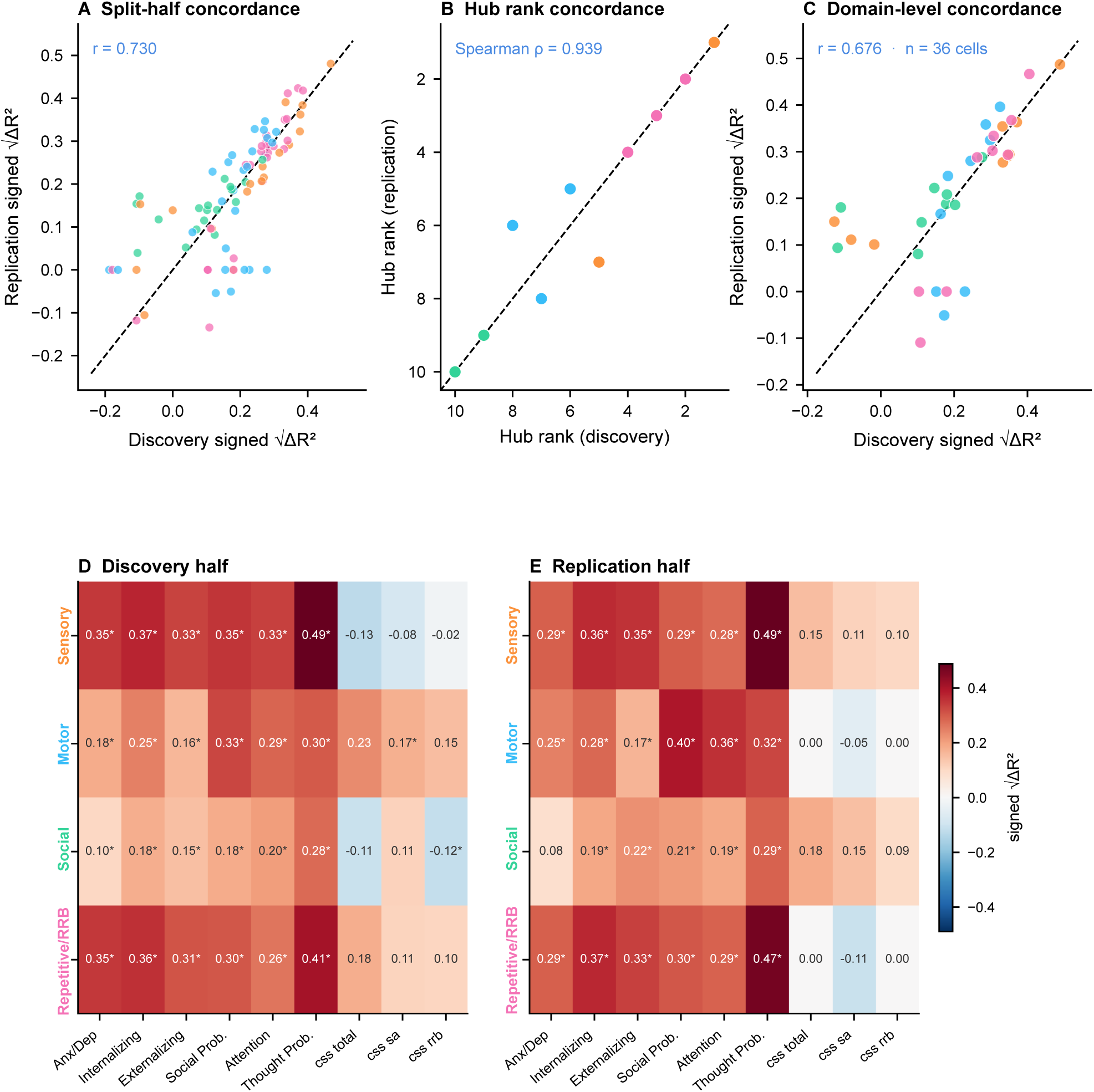
Phenotypic architecture is reproducible across independent SPARK subsets. Internal reproducibility was evaluated by dividing SPARK into independent discovery and replication subsets stratified by sex and age quartile. Signed √ΔR² associations were estimated independently in each subset, with hubness and FDR correction calculated separately within each half. Colors denote behavioral domains. **(A)** Split-half concordance across the 90 predictor × outcome associations. Each point represents the signed √ΔR² estimate for the same association in the discovery and replication halves; the dashed identity line indicates identical estimates. Association effects were strongly concordant across subsets (Pearson r =.730). **(B)** Concordance of independently estimated predictor hub ranks. Each point represents one of the 10 behavioral predictors; the dashed identity line denotes identical rank in the two subsets. Hub ordering was highly reproducible (Spearman ρ =.939). **(C)** Domain-level split-half concordance. Predictor effects were aggregated into Sensory, Motor, Social, and Repetitive/RRB domains, yielding 36 domain × outcome cells. Domain-level signed √ΔR² estimates were concordant between discovery and replication subsets (Pearson r =.676; n = 36 cells). The dashed identity line indicates identical estimates. **(D–E)** Domain-level signed √ΔR² architecture estimated independently in the discovery (D) and replication (E) halves. Rows represent Sensory, Motor, Social, and Repetitive/RRB domains and columns represent the six CBCL psychopathology and three ADOS CSS outcomes. Cell values give signed √ΔR²; asterisks indicate associations surviving FDR correction within the corresponding subset. Color indicates the direction and magnitude of the association.

Participants were randomly divided into discovery and replication halves stratified by sex and age quartile and signed √ΔR² was estimated independently within each subset for all 90 predictor–outcome combinations. Discovery and replication estimates showed substantial concordance (Pearson *r* =.730, *p* <.001; **Figure 3A**). Larger effects, particularly those involving sensory-repetitive and motor predictors with CBCL outcomes, were recovered consistently across subsets, whereas smaller effects showed greater sampling variability. Thus, the overall cross-domain effect-size structure was independently recoverable from two nonoverlapping subsets of SPARK. The higher-order organization of the architecture was even more stable. Hubness was recalculated independently within each half using each subset’s own FDR-corrected associations [30–33]. Predictor ranks were highly concordant (Spearman ρ =.939, *p* <.001; **Figure 3B**). RBS-R Sensory, Obsessive, Ritualistic, and Stereotyped occupied ranks 1–4 in both subsets, whereas SCQ Social and SCQ Communication remained ranks 9 and 10. Rank movement was confined to the middle tier. The stronger agreement in predictor ranks (ρ =.939) than in individual effects (*r* =.730) indicates that modest sampling variation in individual associations had comparatively little influence on the higher-order organization of the phenotype.

I next examined whether reproducibility was retained after aggregation into broader behavioral domains. The predictor-level effects were summarized into Sensory, Motor, Social, and Repetitive/RRB domains, yielding 36 domain × outcome associations.

Discovery and replication estimates remained concordant at this level (Pearson *r* =.676; *n* = 36 cells; **Figure 3C**). Likewise, independently estimated domain-level fingerprint matrices showed similar organization across the discovery and replication subsets (**Figure 3D–E**), with stronger coupling to CBCL psychopathology than to ADOS CSS outcomes and preservation of the dominant sensory/repetitive architecture. Together, these analyses show that the phenotypic architecture is reproducible at three complementary levels: individual predictor-outcome effects, predictor hub ordering, and aggregated domain-level association structure. Although individual effect estimates varied between subsets, the overall hub hierarchy remained highly stable, supporting robustness of the SPARK architecture.

### Phenotypic Architecture Persists but Reorganizes After Anxiety Adjustment

Shared anxiety-related variance could contribute to the sensory-repetitive centrality observed in the primary architecture [17, 34, 35]. I therefore re-estimated predictor-outcome effects after adding CBCL Anxious/Depressed to the baseline covariates and compared the resulting effect-size and hubness profiles with the corresponding baseline architecture (**Figure 4**). Anxiety adjustment broadly attenuated cross-domain effects (**Figure 4A–B**), but the degree of attenuation varied across predictor-outcome combinations. Some of the largest reductions were observed for associations with CBCL Internalizing Problems, consistent with its overlap with Anxious/Depressed scale. RBS-R Sensory × Internalizing decreased from signed √ΔR² =.387 to.126 (change in absolute effect magnitude = −.261), and RBS-R Ritualistic × Internalizing decreased from.344 to

**Figure 4.**
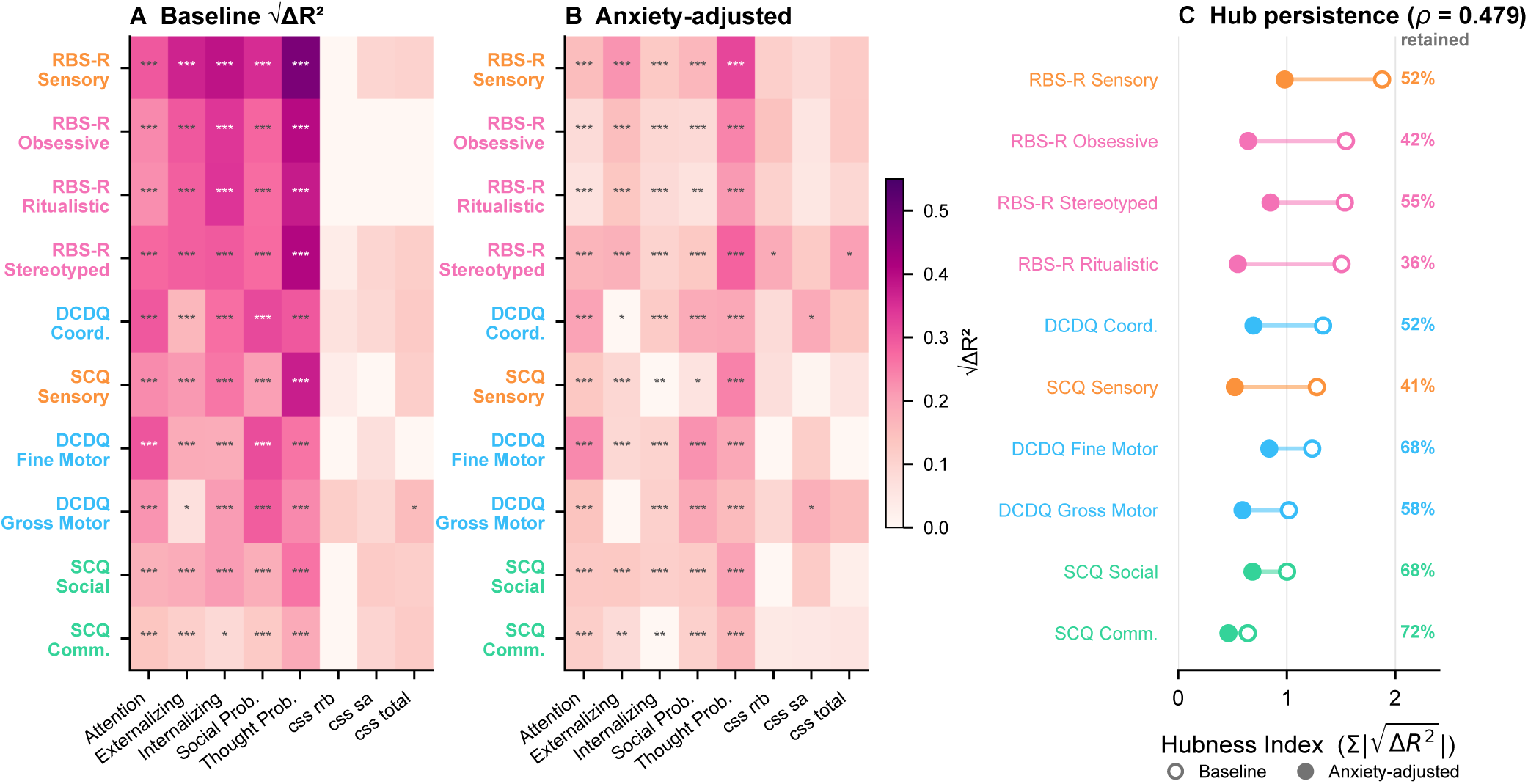
Phenotypic architecture persists but reorganizes after adjustment for anxiety-related variance. The SPARK phenotypic architecture was re-estimated after inclusion of CBCL Anxious/Depressed as an additional covariate. Baseline and anxiety-adjusted analyses used the same matched outcome set, with FDR correction performed separately for each model. **(A)** Baseline signed √ΔR² architecture for the outcomes included in the anxiety-adjustment analysis. **(B)** Corresponding architecture after adjustment for CBCL Anxious/Depressed. Adjustment broadly attenuated effect magnitudes, particularly for associations involving Internalizing, while substantial cross-domain associations remained. **(C)** Change in predictor hub organization following anxiety adjustment. Connected estimates show the baseline and adjusted position of each predictor, illustrating both persistence and reorganization of the hierarchy. Sensory-repetitive dimensions remained prominent after adjustment, although their relative ordering changed.

.084 (−.260). In contrast, substantial associations with other psychopathology outcomes remained after adjustment. RBS-R Sensory × Thought Problems, for example, decreased from.478 to.319 but remained among the largest effects in the adjusted architecture.

Anxiety adjustment therefore removed substantial shared variance without uniformly eliminating cross-domain associations. The predictor hierarchy also reorganized after adjustment (**Figure 4C**). Using matched outcome sets and model-specific FDR correction, baseline and anxiety-adjusted hub ranks showed moderate concordance (Spearman ρ =.479). RBS-R Stereotyped and Sensory scales occupied the top two adjusted positions, while DCDQ Coordination and Fine Motor rose to ranks 3 and 4. RBS-R Obsessive and Ritualistic scales, in contrast, moved from ranks 2 and 4 at baseline to ranks 7 and 8 after adjustment, and SCQ Sensory shifted from rank 6 to rank 9. SCQ Communication remained rank 10.

These findings indicate that anxiety-related variance contributes meaningfully to both the magnitude and relative organization of cross-domain associations in SPARK. However, adjustment did not eliminate the broader architecture: substantial associations remained, and sensory-repetitive dimensions continued to occupy prominent positions within the hierarchy. The primary architecture therefore cannot be reduced simply to covariance with anxiety, although its finer organization is sensitive to anxiety adjustment.

### Phenotypic Architecture Reorganizes Across Development

I next asked whether the relative organization of the phenotypic architecture was stable across development. Hubness was estimated independently within four SPARK age bands (4–8, 9–12, 13–17, and ≥18 years), and age-specific predictor ranks were compared to quantify developmental stability (**Figure 5**).

**Figure 5.**
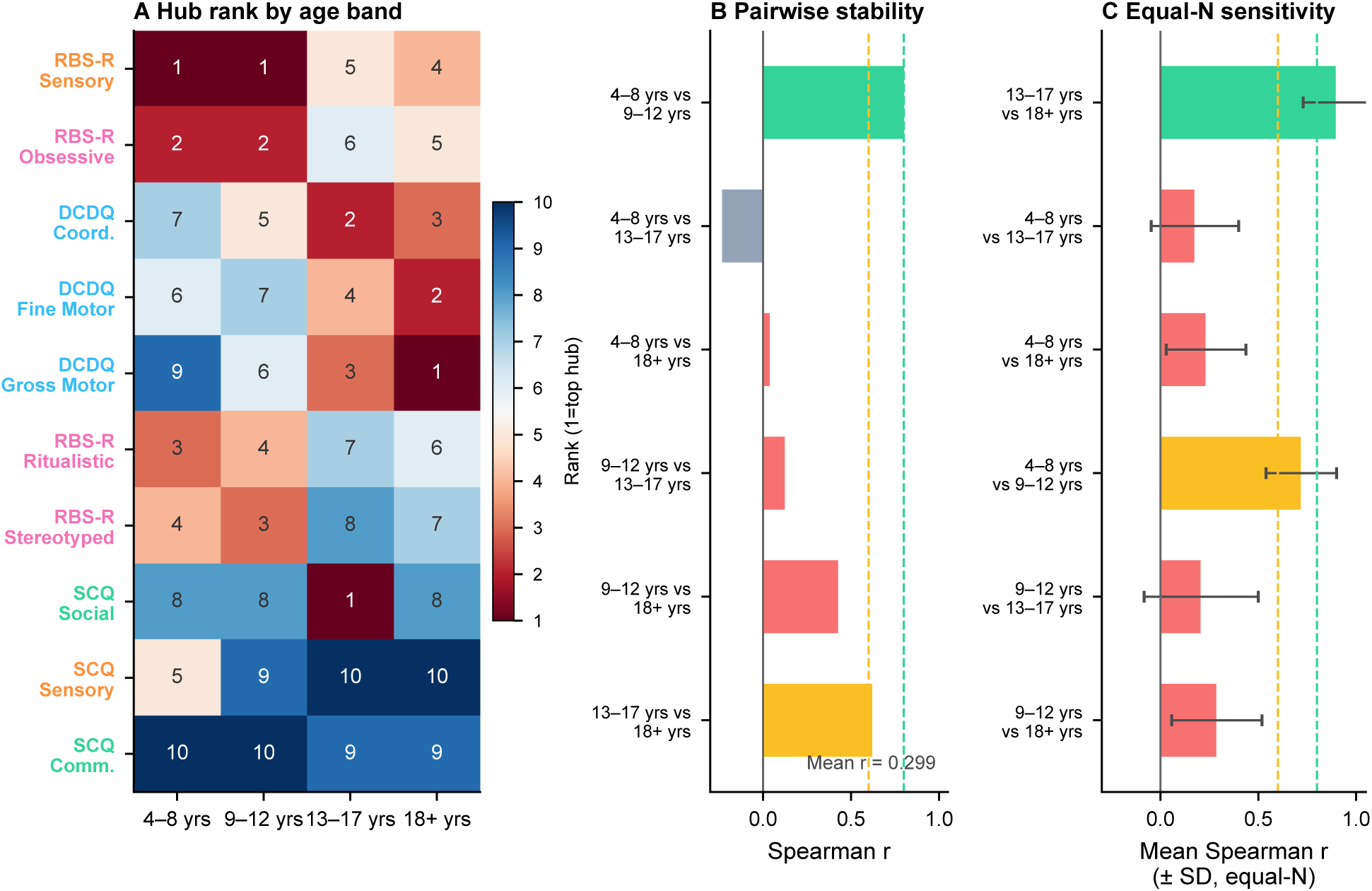
Phenotypic hub architecture reorganizes across development. Hubness was estimated independently within four SPARK age bands (4–8, 9–12, 13–17, and ≥18 years), with sex retained as a covariate. Developmental stability was evaluated from concordance of independently estimated hub rankings. **(A)** Age-specific predictor hub ranks. Sensory-repetitive dimensions dominated the childhood hierarchy, whereas motor dimensions rose markedly in adolescence and adulthood. **(B)** Pairwise Spearman concordance between age-specific hub rankings. Rank organization was similar between the two childhood groups (ρ =.806, *p* =.005), substantially less concordant across the childhood-to-adolescent transition, and more similar again between adolescent and adult architectures (ρ =.624, *p* =.054). **(C)** Equal-*N* sensitivity analysis. Age groups were repeatedly subsampled to equal numbers of observations (*n* = 17,866; 20 iterations), and hub-rank concordance was recomputed. Points and error bars represent mean Spearman ρ ± SD across iterations. The developmental pattern remained after equalizing sample size, arguing against unequal age-group sample sizes as the primary explanation for the observed reorganization.

The hub hierarchy changed substantially across age groups (**Figure 5A**). Sensory-repetitive dimensions occupied the highest positions during childhood: RBS-R Sensory and Obsessive scale ranked first and second in both the 4–8-and 9–12 age groups. By adolescence, however, motor dimensions had risen sharply in relative rank. DCDQ Coordination, Gross Motor, and Fine Motor occupied ranks 1–3 at ages 13–17, and DCDQ Gross Motor, Fine Motor, and Coordination occupied the top three positions in adults.

DCDQ Gross Motor showed one of the largest shifts, moving from near the bottom of the childhood hierarchy to the highest-ranked position in adulthood. Pairwise rank correlations quantified this developmental pattern (**Figure 5B**). Hub ordering was strongly concordant between the two childhood groups (Spearman ρ =.806, *p* =.005) but showed little correspondence across several comparisons spanning the childhood-to-adolescent transition. The 4–8 versus 13–17 comparison was negative (ρ = −.236), whereas the adolescent and adult architectures were more similar (ρ =.624, *p* =.054). These results suggest relative stability within childhood, followed by a substantial reorganization around adolescence and partial stabilization thereafter.

Because sample size differed across age groups, I repeated the analysis after equalizing the number of observations (**Figure 5C**). Each age group was repeatedly subsampled to *n* = 17,866 observations across 20 iterations before hub ranks were recomputed. The same broad pattern remained: within-developmental-period comparisons showed substantially greater rank concordance than comparisons spanning the childhood-to-adolescent boundary. The developmental reorganization therefore was not explained simply by unequal sample sizes across age groups. Together, these analyses identify age as an important axis of heterogeneity in the SPARK architecture. Sensory-repetitive dimensions dominate the relative hierarchy during childhood, whereas motor dimensions become increasingly prominent during adolescence and adulthood. This analysis concerns relative hub organization; whether the absolute strength of domain-psychopathology coupling follows the same developmental pattern is addressed separately below. Although not necessarily reflecting changes in the absolute strength of associations with psychopathology, these age-related shifts reflect changes in the relative organization of the hub hierarchy.

### Phenotypic Architecture Replicates in the Simons Simplex Collection

Having established the architecture and its internal reproducibility in SPARK, I next tested whether its major features generalized to the independent Simons Simplex Collection (SSC). Because SSC provided the six CBCL psychopathology outcomes but not the three ADOS CSS outcomes used in the full SPARK architecture, cross-cohort comparisons were restricted to the common CBCL outcome space (**Figure 6**).

**Figure 6.**
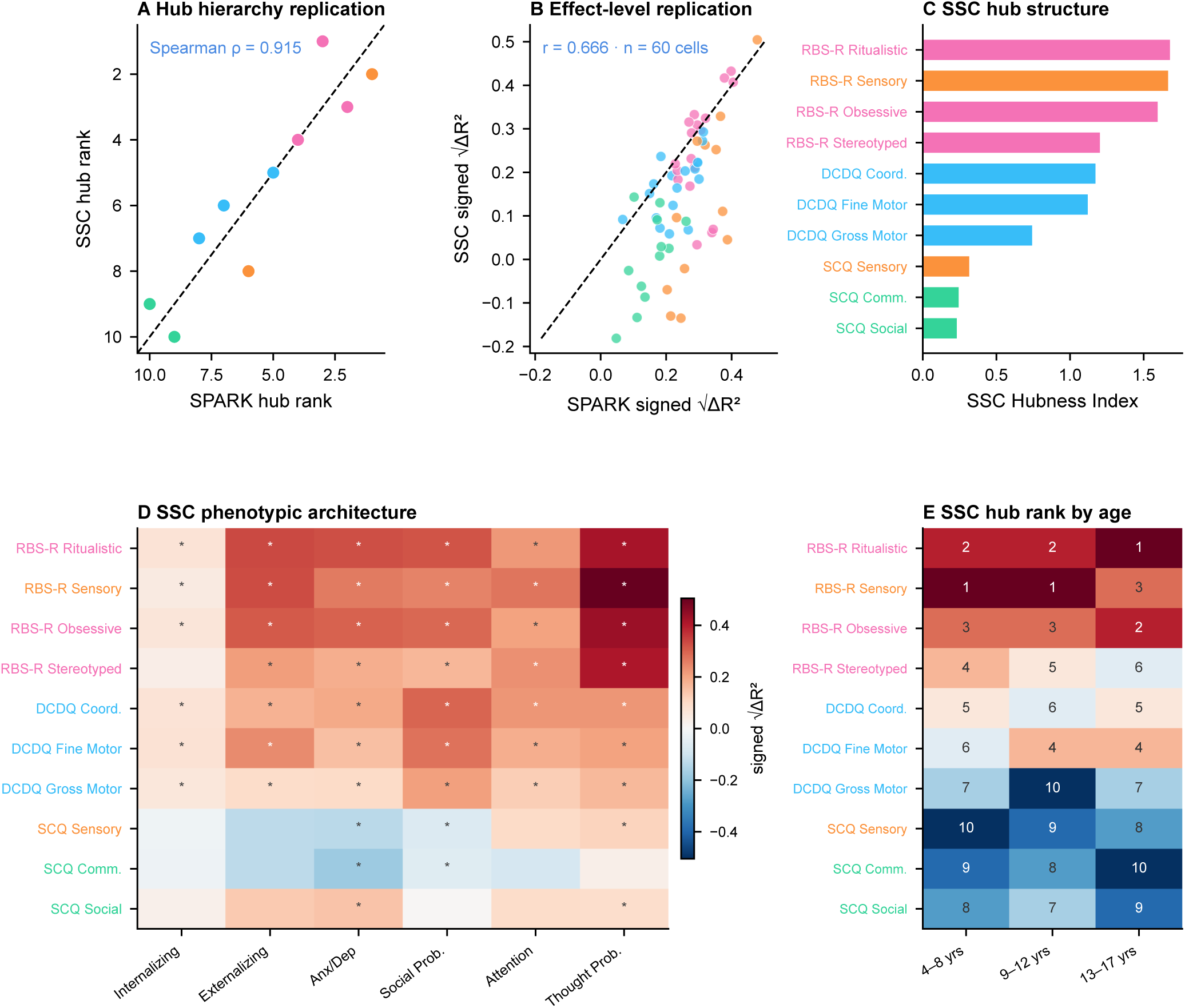
Phenotypic architecture replicates in the independent Simons Simplex Collection. External replication was evaluated in the Simons Simplex Collection (SSC) using the six CBCL psychopathology outcomes available in both SPARK and SSC. Analyses used the same signed √ΔR² framework, permitting comparison of both individual associations and higher-order hub organization across cohorts. **(A)** Cross-cohort concordance of predictor hub ranks. SPARK and SSC showed strong preservation of the hub hierarchy (Spearman ρ =.915, *p* <.001). The four RBS-R dimensions occupied the four highest positions in both cohorts, and DCDQ Coordination ranked fifth in both. **(B)** Cross-cohort concordance of the 60 predictor × CBCL associations available in both datasets. Signed √ΔR² estimates were correlated across cohorts (Pearson *r* =.666, *p* <.001), demonstrating replication at the individual-association level in addition to the hub hierarchy. **(C)** SSC Hubness Index across the 10 behavioral predictors. RBS-R Ritualistic, Sensory, Obsessive, and Stereotyped occupied the four highest positions. **(D)** SSC signed √ΔR² fingerprint across the six CBCL outcomes. Color represents association magnitude and significance markers denote associations surviving FDR correction. **(E)** Age-specific SSC hub ranks across ages 4–8, 9–12, and 13–17 years. Rank organization was highly stable across the three age bands (pairwise Spearman ρ =.842–.891; mean ρ =.871), with RBS-R Sensory, Ritualistic, and Obsessive remaining within the top three positions. Thus, SSC strongly replicated the overall sensory-repetitive-centered architecture but showed greater developmental stability than the pronounced motor-rank ascent observed in SPARK.

The higher-order predictor hierarchy replicated strongly across cohorts (**Figure 6A**). SPARK and SSC hub ranks were highly concordant (Spearman ρ =.915, *p* <.001). All four RBS-R dimensions occupied the top four positions in both cohorts, and DCDQ Coordination ranked fifth in both. The exact ordering within the sensory-repetitive cluster differed slightly: RBS-R Sensory ranked first in SPARK, whereas RBS-R Ritualistic ranked first in SSC. Nevertheless, the dominant organization was preserved, with sensory-repetitive dimensions concentrated at the top and social-communication dimensions generally occupying lower positions. Replication was also evident at the individual-association level (**Figure 6B**). Across the 60 predictor × CBCL associations available in both cohorts, signed √ΔR² estimates were positively correlated (Pearson *r* =.666, *p* <.001). Thus, external replication was not limited to the rank ordering produced by the Hubness Index; the underlying pattern of individual predictor–psychopathology associations was also preserved across cohorts.

The SSC architecture itself showed a pronounced sensory-repetitive hierarchy (**Figure 6C–D**). RBS-R Ritualistic scale had the highest Hubness Index (H = 1.68), followed by RBS-R Sensory (H = 1.67), Obsessive (H = 1.59), and Stereotyped scale (H = 1.20). DCDQ Coordination ranked fifth (H = 1.17), followed by Fine Motor and Gross Motor, whereas the SCQ dimensions occupied the lower portion of the hierarchy. The SSC effect-size fingerprint similarly showed broad sensory-repetitive and motor associations with CBCL psychopathology, providing a direct view of the association structure underlying the replicated hub ordering.

I also examined the SSC hierarchy across the three comparable age bands available in that cohort (4–8, 9–12, and 13–17 years; **Figure 6E**). In contrast to the stronger developmental reorganization observed in SPARK, SSC hub rankings remained highly concordant across age groups: Spearman ρ =.891 for 4–8 versus 9–12 years,.879 for 4–8 versus 13–17 years, and.842 for 9–12 versus 13–17 years (mean pairwise ρ =.871). RBS-R Sensory, Ritualistic, and Obsessive remained within the top three positions across all three age bands. SSC therefore provides strong external replication of the overall sensory-repetitive-centered architecture, while placing an important boundary on the developmental finding from SPARK. The broad hub hierarchy and individual association structure generalize across cohorts, but the pronounced childhood-to-adolescent motor-rank ascent observed in SPARK was not reproduced in SSC. Developmental reorganization of the finer predictor hierarchy should therefore be considered cohort-or context-sensitive pending further replication.

### Developmental Changes in Domain–Psychopathology Coupling

The age-stratified hub analysis showed that the relative organization of the SPARK architecture changes across development. I next asked a complementary question: whether the absolute strength of coupling between broad behavioral domains and psychopathology also varies developmentally. Behavioral measures were aggregated into Sensory-Repetitive (SR), Motor, and Social domain composites, and their associations with a CBCL psychopathology composite were examined across four developmental bands (0–4, 4–8, 8–12, and 12–18 years; **Figure 7**). Motor effects were not estimable at ages 0–4 because DCDQ measures were unavailable in that age range. All 11 estimable domain × age-band associations survived FDR correction (*q* <.001). SR scale showed the strongest population-level psychopathology coupling across developmental bands (**Figure 7A**). The largest effect occurred at ages 0–4 (signed √ΔR² ≈.50, *n* = 4,132), followed by lower effects at 4–8 (.431, *n* = 7,998) and 8–12 (.439, *n* = 6,084), with a similarly lower estimate at 12–18 years. Pairwise comparisons supported an early-childhood peak, whereas the continuous linear age trend was not significant. The pattern therefore indicates particularly strong SR–psychopathology coupling in early childhood rather than a simple monotonic decline across development. Motor–psychopathology coupling was intermediate and comparatively stable across the age ranges for which DCDQ data were available, whereas Social coupling was consistently weakest. Bayesian estimates supported the same domain ordering: relative to SR, coupling was lower for Motor (β = −.230, 95% CrI [−.265, −.194]) and Social (β = −.353, 95% CrI [−.437, −.269]), with *P*(direction) = 1.00 for both contrasts. Thus, the developmental rise of motor dimensions in relative hub rank in **Figure 5** does not imply that motor features become more strongly coupled to psychopathology than sensory-repetitive features.

**Figure 7.**
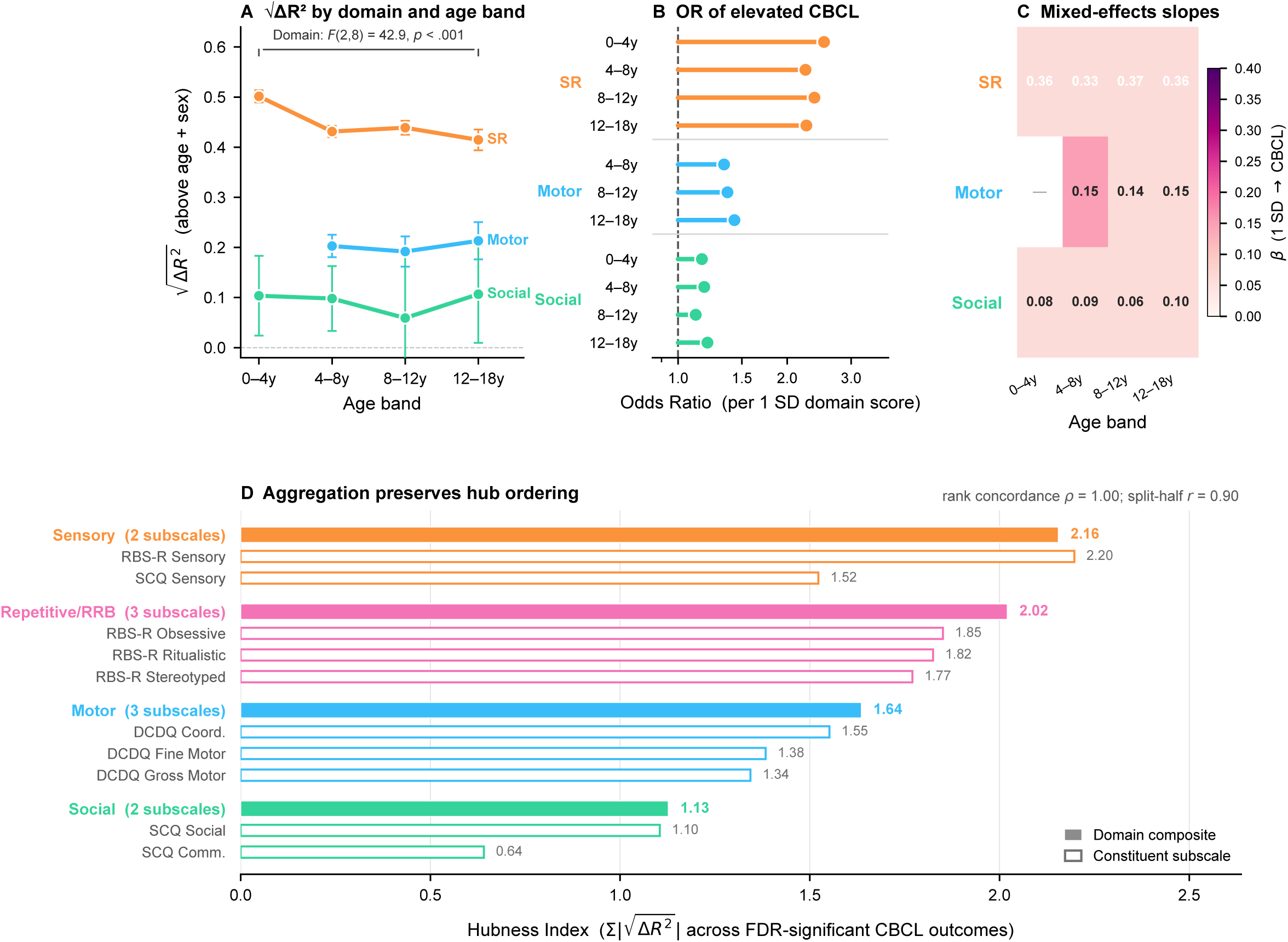
Developmental patterning of domain–psychopathology coupling. Behavioral measures were aggregated into Sensory-Repetitive (SR), Motor, and Social domains to examine developmental variation in their association with psychopathology. Analyses were conducted across developmental bands spanning ages 0–18 years; Motor estimates were unavailable at ages 0–4 because DCDQ measures were not available in that age range. **(A)** Population-level signed √ΔR² between each behavioral domain and the CBCL psychopathology composite across developmental bands. SR showed the strongest coupling throughout development, with its largest effect during early childhood. Motor coupling was intermediate and comparatively stable across the age ranges in which it was available, whereas Social coupling was consistently weaker. **(B)** Association between behavioral domain scores and clinically elevated psychopathology (CBCL T ≥ 63) across developmental bands, expressed as odds ratios with confidence intervals. Estimates quantify the change in odds of clinically elevated psychopathology associated with increasing domain score within each developmental band. **(C)** Individual-level domain–psychopathology slopes across developmental bands estimated from mixed-effects models. SR slopes were consistently largest, Motor slopes intermediate, and Social slopes smaller, complementing the population-level effect-size pattern in panel A. **(D)** Validation of domain aggregation. Filled bars represent hubness of the aggregated domain composites, and outlined bars show the corresponding constituent behavioral measures, demonstrating preservation of the broader ordering of the phenotypic architecture after aggregation.

The clinical-threshold analysis provided a complementary measure of this domain ordering (**Figure 7B**). Higher SR scores were associated with the greatest odds of clinically elevated psychopathology (CBCL T ≥ 63) across developmental bands, with Motor and Social associations generally smaller. This analysis therefore translated the continuous domain-psychopathology relationship into a clinically interpretable outcome while preserving the same broad ordering observed for signed √ΔR². Individual-level slopes likewise showed persistent domain differentiation (**Figure 7C**). In the stacked population model (*n* = 62,846 observations), SR slopes were large and comparatively stable across developmental bands (β =.356,.332,.371, and.363). Motor slopes were smaller (approximately β =.14-.15), whereas Social slopes were weaker and more variable (β =.057-.101). A significant score × Domain × Band interaction (*F*(6, 62,826) = 2.6, *p* =.017) nevertheless indicated that developmental variation differed across domains. It is worth highlighting that while at population level, SR explains less variation in psychopathology after early childhood, at the individual level, the relationship between higher SR scores and higher psychopathology scores persist the same across age groups.

Finally, I evaluated whether aggregation into broad domains preserved the organization of the underlying phenotypic architecture (**Figure 7D**). Aggregating the individual measures into broader domains preserved their overall ordering, supporting the use of domain composites in the developmental analyses, and indicating that the developmental domain results are not simply an artifact of collapsing heterogeneous behavioral measures into composite scores. Together, the developmental analyses distinguish relative architectural organization from absolute psychopathology coupling.

Motor dimensions rise substantially in relative hub rank across the SPARK age-stratified architecture, but sensory-repetitive features remain the behavioral domain most strongly coupled to psychopathology throughout childhood and adolescence. The particularly strong population-level SR coupling in early childhood, alongside comparatively stable individual-level slopes, further indicates that developmental changes depend on the level of phenotypic organization being considered.

## Discussion

ASD is defined diagnostically by a particular set of behavioral features [36], but diagnosis does not necessarily reveal how those features are organized within the broader phenotype. Across complementary analyses in SPARK and independent replication in SSC, I identified a reproducible cross-domain architecture in which sensory-repetitive features occupied disproportionately central positions. This organization emerged across individual associations, hub rankings, and multivariate structure, and was reproducible both within SPARK and in the independent SSC cohort. Yet centrality varied across age and did not necessarily track the strength of associations with psychopathology. Anxiety adjustment altered the magnitude and ordering of associations without eliminating the broader sensory-repetitive-centered structure, and developmental analyses showed that a feature’s relative position within the architecture can change without a corresponding change in its association with psychopathology. Together, these findings suggest that autism-related heterogeneity is not simply multidimensional: it is organized, and that organization provides information that diagnostic prominence and severity alone do not capture.

### What defines autism is not necessarily what organizes its broader phenotype

Here, the central conceptual finding is a dissociation between diagnostic prominence and architectural centrality. Diagnostic criteria identify features that characterize autism, and severity measures quantify the extent to which those features are expressed [10, 37]. Architectural centrality asks a different question: which behavioral features are most broadly and strongly connected to variation elsewhere in the phenotype? [11]. The study shows that sensory-repetitive dimensions occupied the top of the hub hierarchy, whereas SCQ Communication occupied its lowest position. The same asymmetry emerged from an unsupervised decomposition of the association matrix: the dominant principal component accounted for 54.8% of variance and separated strongly connected sensory-repetitive features from comparatively peripheral social-communication features. Moreover, this architecture was expressed primarily through associations with psychopathology. Fifty-nine of 60 predictor–CBCL associations survived FDR correction, compared with only one of 30 associations with clinician-rated ADOS CSS. Thus, the behavioral dimensions most deeply embedded in the broader phenotype were not simply those showing the strongest relationships with clinician-rated autism severity.

This distinction extends existing efforts to move beyond a unitary conception of autism severity [37, 38]. Autism phenotypes are multidimensional, and previous factor-analytic and genetic work has shown that diagnostic features themselves resolve into separable dimensions rather than a single severity axis [38, 39]. Recent large-scale studies have similarly demonstrated that phenotypic heterogeneity can reveal biologically meaningful structure not apparent from diagnostic categories alone [39–41]. This study adds a complementary level of organization: even after dimensions have been separated, they differ in how strongly they are connected to the rest of the phenotype (**Figure 2**). A feature can therefore be diagnostically important without being architecturally central, or architecturally central without being the feature most characteristic of diagnosis. Social-communication differences are fundamental to the clinical definition of autism [36], yet sensory, repetitive, motor, and psychiatric features often account for substantial variation in everyday functioning and clinical complexity [37, 42, 43]. Architectural centrality does not challenge the diagnostic importance of social communication. Rather, it suggests that the features most useful for identifying a condition need not be the same features most connected to variation within that condition.

### Sensory-repetitive centrality is a reproducible property of the architecture

An architectural account is useful only if the inferred organization is sufficiently stable to survive changes in sample and analytic representation [44, 45]. The sensory-repetitive-centered pattern proved robust across several levels of analysis. Across independent SPARK halves, predictor-level effects were substantially concordant, hub rankings were even more stable, and the broader domain-level architecture was consistently preserved (**Figure 3**). Most importantly, the higher-order hierarchy generalized to SSC: SPARK and SSC hub ranks correlated at ρ =.915 (**Figure 6**), suggesting that sensory-repetitive centrality is not simply a consequence of a particular summary statistic or a chance configuration of SPARK.

Centrality of sensory-repetitive features is not unexpected. Sensory differences emerge early in development and can influence how children engage with both physical and social environments [14–16]. Repetitive behaviors can similarly shape patterns of exploration and interaction [19]. Features that emerge early and intersect repeatedly with emotional regulation, learning, social experience, and behavior have more opportunities to become associated with multiple downstream domains [20, 46]. In this sense, broad behavioral centrality could arise not because a feature is intrinsically more severe, but because it is involved in more processes through which development unfolds. This interpretation is compatible with mechanistic work implicating sensory integration and sensory-higher-order interactions in autism. Altered organization of the functional connectome hierarchy has been associated with differences in the integration of sensory information into higher-order systems [47], while computational accounts have implicated altered weighting of sensory evidence and prior expectations [48]. Experimental work in animal models further demonstrates that perturbations of peripheral sensory processing can influence social and anxiety-like behaviors [49]. These studies do not establish the mechanism underlying the behavioral architecture identified here, nor does centrality itself imply causal precedence. They do, however, make the broad connectivity of sensory-repetitive behavior biologically plausible and suggest concrete hypotheses for future multimodal and longitudinal studies.

The architecture also complements recent attempts to decompose autism heterogeneity at phenotypic and genetic levels. Large-scale work in SPARK and SSC has identified separable sensory-motor, repetitive, social, and communication dimensions and shown that phenotypic subdivisions can carry distinct genetic information. Rather than asking which dimensions or subgroups exist, the present analysis asks how those dimensions relate to one another and to clinically consequential outcomes. These perspectives are complementary: decomposition identifies the components of heterogeneity, whereas architectural analysis asks how those components are organized.

### Anxiety contributes to the architecture but does not explain it

The strong connection between sensory-repetitive features and psychopathology raises an important alternative explanation. Sensory and repetitive behaviors are closely associated with anxiety [17, 18, 50], and anxiety-related variance is represented across several psychopathology measures [51, 52]. Sensory-repetitive features might therefore appear central because both they and the outcomes reflect a common anxiety-related dimension. The anxiety-adjustment analysis indicates that this explanation is partly correct but insufficient. Adding Anxious/Depressed symptoms to the baseline covariates substantially attenuated many associations and reorganized portions of the hub hierarchy (**Figure 4**). Internalizing associations showed some of the largest reductions, and several repetitive-behavior dimensions moved substantially in rank. Anxiety-related variance is therefore not incidental to the architecture; it contributes meaningfully to both its magnitude and its finer organization.

At the same time, substantial cross-domain structure remained. RBS-R Sensory retained a prominent position, and strong associations with outcomes such as Thought Problems persisted despite adjustment (**Figure 4**). The architecture therefore cannot be reduced to a chain in which sensory-repetitive behavior indexes anxiety and anxiety alone accounts for the remaining cross-domain associations. A more plausible interpretation is that anxiety represents one of several pathways connecting sensory-repetitive features with broader clinical variation, consistent with evidence linking sensory features to anxiety through multiple direct and indirect pathways [16, 17], as well as work implicating intolerance of uncertainty, interoception, autonomic regulation, and sleep in relationships between sensory experience and emotional symptoms [34, 53–55]. These data cannot determine which of these pathways generates the residual architecture after anxiety adjustment; however, anxiety seems to contribute to sensory-repetitive centrality but is not a sufficient explanation for it.

### Development separates architectural position from clinical coupling

Development provides perhaps the clearest demonstration that architectural centrality is not interchangeable with symptom magnitude or clinical importance. In SPARK, hub organization changed substantially across age. Sensory-repetitive dimensions dominated the childhood hierarchy, whereas motor dimensions rose sharply in relative rank during adolescence and adulthood (**Figure 5**). In contrast, SSC hub rankings remained highly stable, with sensory-repetitive dimensions retaining the leading positions (**Figure 6**). Although, the prominence of sensory-repetitive features and comparatively lower position of social-communication dimensions was reproduced across cohorts. This suggests that the architecture may contain both stable and context-sensitive components), and differences in cohort composition, measurement, developmental range, ascertainment, or other contextual factors may influence how that common architecture is expressed at particular ages. Longitudinal replication will be necessary to determine which developmental changes represent general maturational processes and which depend on cohort or context.

The SPARK developmental analyses, nevertheless, reveal an important principle independent of whether the motor-rank transition generalizes universally. Relative architectural position and absolute clinical coupling are separable properties. Although motor features rose dramatically in relative hub rank with age in SPARK (**Figure 5**), sensory-repetitive features remained the domain most strongly associated with psychopathology across developmental bands (**Figure 7**). Motor centrality therefore did not mean that motor difficulties replaced sensory-repetitive features as the strongest correlate of psychiatric burden, suggesting that rank is inherently relational: A behavioral feature can rise in architectural centrality because its relationships strengthen, because relationships involving other features weaken, or because the configuration of the broader phenotype changes around it. None of these possibilities require that the feature becomes the strongest predictor of a particular clinical outcome. A phenotype can reorganize when relationships between individual traits remain clinically meaningful, just as stable symptom magnitude does not necessarily imply stability in the broader phenotypic structure [11, 16, 56].

### Clinical and conceptual implications

Architectural centrality identifies features that may contain disproportionate information about broader clinical variation. Early-childhood results are particularly relevant (**Figure 5**), as sensory and repetitive behaviors are already observable at this stage, while psychiatric presentations may still be developing. This raises the possibility that sensory-repetitive phenotyping could contribute to early identification of children who warrant closer assessment of emerging emotional and behavioral difficulties. This possibility should be tested directly in longitudinal cohorts. More broadly, architectural analysis offers a different way of approaching heterogeneity. Much autism research has focused on identifying subgroups, latent dimensions, trajectories, or biological correlates [22, 39–41], and focus on the forms of heterogeneity that exist and what explains them. In contrast, architectural centrality asks how heterogeneity itself is organized [11], which could provide a useful behavioral bridge to biological work. Network, genetic, and computational analyses increasingly characterize autism in terms of distributed systems rather than isolated deficits [47, 57]. Behavioral phenotypes can likewise be viewed not as interchangeable expressions of a single severity dimension, but as interconnected components whose relationships are themselves informative. The present findings establish that such relational structure can be measured reproducibly at the behavioral level. Determining whether behavioral hubs correspond to particular genetic, neural, or computational architectures is now an empirical question rather than an assumption.

### Limitations and future directions

Several limitations define the appropriate scope of these conclusions. First, the developmental analyses are primarily cross-sectional. Age-stratified differences characterize how phenotypic architecture differs among individuals observed at different developmental stages; they do not demonstrate within-person reorganization. This distinction is especially important given the difference between SPARK and SSC developmental patterns. Longitudinal measurement of the same behavioral domains will be necessary to determine whether individuals undergo systematic architectural transitions and whether such transitions predict later functioning or psychopathology.

Second, several measures are parent-reported, creating potential shared-method variance. This is particularly relevant to the strong associations between behavioral predictors and CBCL psychopathology. The much weaker associations with clinician-rated ADOS CSS demonstrate that the architecture is not uniform across measurement modalities, but they do not by themselves distinguish substantive outcome differences from informant or instrument effects. Replication using clinician-administered, performance-based, ecological, and self-report measures will be important.

Third, centrality is an associational property. A behavioral hub is broadly connected within the measured phenotype, but this does not establish that it causally generates downstream features. The mechanistic literature discussed above motivates hypotheses about sensory processing, repetitive behavior, predictive control, and developmental cascades, but causal direction cannot be inferred from the present analyses. Longitudinal, genetically informed, intervention, and multimodal studies will be required to determine whether highly central behavioral features are causes, consequences, markers of shared mechanisms, or some combination of these.

Fourth, architecture necessarily depends on what is measured. The present study sampled motor, sensory-repetitive, social-communication, psychopathology, and clinician-rated autism dimensions using measures available at scale in SPARK and SSC. Other domains (adaptive functioning, language, sleep, attention, executive function, interoception, and daily living skills) could alter the estimated architecture if incorporated comprehensively. Hubness should therefore be interpreted as centrality within a defined phenotypic measurement space rather than an intrinsic and context-free property of a behavior.

Finally, the contrast between strong cross-cohort replication of the overall architecture and weaker replication of its developmental configuration highlights the importance of separating robust organizing principles from cohort-specific detail. SPARK and SSC differed in sample composition, available outcomes, age coverage, and measurement structure. Rather than treating those differences only as limitations, future work could use multiple cohorts deliberately to identify which components of phenotypic architecture remain invariant across populations and which respond to developmental or environmental context.

## Conclusion

Autism-related heterogeneity is often described by cataloguing the many ways autistic individuals differ. This study suggests that the relationships among those differences are themselves informative. Across SPARK and SSC, sensory-repetitive features occupied a reproducible central position in the broader behavioral phenotype, despite not simply mirroring clinician-rated autism severity. Their centrality was partly–but not wholly–related to anxiety, and their strong association with psychopathology persisted across development. At the same time, the study shows why centrality should not be mistaken for severity, diagnostic importance, or clinical risk. The global architecture was reproducible across cohorts, whereas its developmental configuration was more variable; within SPARK, motor features rose in relative architectural position without replacing sensory-repetitive features as the domain most strongly coupled to psychopathology. What defines autism, what organizes its phenotype, and what carries clinical risk are therefore overlapping but distinct properties.

Treating the autistic phenotype as an architecture rather than a catalogue of symptoms makes those distinctions visible. It provides a framework in which heterogeneity is not simply something to partition or reduce, but something whose organization can itself be measured, replicated, and ultimately connected to development and mechanism.

## Acknowledgements

I am grateful to all participants, families research teams in SSC and SPARK as well as SFARI Base staff, this study was only possible due of their efforts.

## STAR Methods

### Resource availability

#### Lead contact

Further information and requests for resources should be directed to and will be fulfilled by the lead contact at.

#### Materials availability

This study did not generate new materials.

#### Data and code availability

De-identified phenotypic data from SPARK and Simons Simplex Collection (SSC) are available through SFARI Base subject to applicable data-use agreements. Analysis code is deposited at [https://github.com/fdzlab-UTRGV/ASD_DevDynamics].

### Experimental model and study participant details

#### Study cohorts

The primary analyses used phenotypic data from SPARK (Simons Foundation Powering Autism Research for Knowledge), a large research cohort of autistic individuals and their families. Analyses were conducted using available behavioral, psychopathology, autism-severity, demographic, and cognitive measures. Because instruments were administered to overlapping but non-identical subsets of participants, analytic sample size varied across analyses and predictor–outcome combinations.

The Simons Simplex Collection (SSC) was used as an independent replication cohort. Replication analyses were restricted to behavioral predictors and outcomes with corresponding measures available in both cohorts. Because the three ADOS calibrated severity score outcomes included in the primary SPARK architecture were not available in the corresponding SSC replication matrix, direct SPARK–SSC effect-size comparisons were restricted to the six shared CBCL outcomes.

### Behavioral measures

#### Motor behavior

Motor coordination was assessed using the Developmental Coordination Disorder Questionnaire (DCDQ). Gross Motor, Fine Motor, and Coordination subscales were included as separate predictors in the primary architectural analyses.

#### Sensory and repetitive behavior

Repetitive behavior was assessed using the Repetitive Behavior Scale-Revised (RBS-R). Sensory, Obsessive, Ritualistic, and Stereotyped subscales were included as separate predictors in the primary analyses.

#### Social communication

Social-communication features were assessed using the Social Communication Questionnaire (SCQ). Social, Communication, and Sensory subscales were included as separate predictors.

Together, these measures yielded 10 behavioral predictors spanning motor, sensory/repetitive, and social-communication features.

#### Psychopathology

Co-occurring psychopathology was assessed using six Child Behavior Checklist (CBCL) scales: Anxious/Depressed, Internalizing, Externalizing, Social Problems, Attention Problems, and Thought Problems.

#### Autism symptom severity

Clinician-rated autism symptom severity was represented by three ADOS calibrated severity scores (CSS): Total, Social Affect, and Restricted/Repetitive Behavior. The primary SPARK architecture therefore comprised 10 behavioral predictors × 9 outcomes, yielding 90 predictor-outcome associations.

### Quantification and Statistical Analysis

#### Covariate-adjusted cross-domain architecture

Cross-domain associations were quantified using the incremental variance explained by each behavioral predictor beyond the prespecified covariates age, sex, and nonverbal IQ (NVIQ). Analyses were conducted separately for each predictor-outcome combination using available observations for the variables required for that model; consequently, analytic sample size varied across cells.

For each predictor XX and outcome YY, two ordinary least-squares regression models were estimated:

### Base model

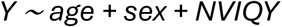

### Full model

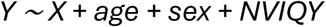

The predictor’s incremental contribution was calculated as:

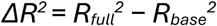

and expressed as a signed square-root effect size:

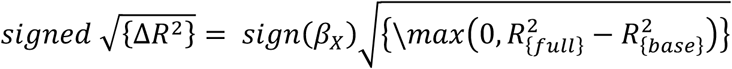

Here, *βX* is the regression coefficient for the behavioral predictor in the full model. The magnitude of signed √ΔR² therefore represents the incremental variance attributable to the predictor beyond the covariates, expressed in correlation-like units, whereas its sign represents the direction of the adjusted predictor–outcome association. Negative values consequently indicate inverse adjusted associations and do not indicate negative explained variance. Statistical significance for each predictor was derived from the corresponding regression coefficient in the full model. Multiple comparisons were controlled using the Benjamini–Hochberg false-discovery-rate (FDR) procedure across the 90 finite predictor–outcome tests. Cells with unavailable estimates were excluded from the number of tests used for FDR correction. This covariate-adjusted signed √ΔR² matrix constituted the primary representation of phenotypic architecture used throughout the study.

#### Hubness Index

To quantify the extent to which individual behavioral features were broadly connected across the phenotype, I defined a Hubness Index for each predictor *i*:

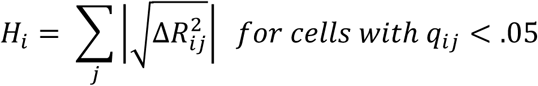

Thus, hubness integrates both the extent of significant associations across outcomes and their magnitude [32, 58, 59]. Predictors with numerous large FDR-significant associations receive higher hubness values than predictors with fewer or weaker associations.

Predictors were ranked in descending order of Hubness Index. Hubness is used here as a descriptive measure of cross-domain architectural centrality within the measured predictor-outcome space and not as causal influence or graph-theory centrality.

#### Principal-component analysis

To determine whether the predictor organization identified by hubness was also evident without explicitly defining a centrality metric, principal-component analysis (PCA) was applied to the 10 × 9 signed √ΔR² matrix. Outcome columns were centered and scaled before decomposition. PCA was performed using singular-value decomposition, and the proportion of matrix variance explained by each component was calculated from its singular values [60]. Predictor scores on the first two components were used to characterize the dominant multivariate organization of cross-domain association profiles.

### Internal reproducibility within SPARK

#### Predictor-level split-half reproducibility

Internal reproducibility was evaluated by dividing SPARK participants into nonoverlapping discovery and replication subsets stratified by sex and age quartile. The complete signed √ΔR² analysis, including model estimation and FDR correction, was performed independently within each subset. Effect-size reproducibility was quantified as the *Pearson* correlation between discovery-and replication-half signed √ΔR² estimates across the 90 predictor × outcome cells. Hubness was then calculated independently in each subset using that subset’s own FDR-corrected associations. Preservation of predictor ordering was quantified using Spearman rank correlation across the 10 discovery-and replication-half hub ranks.

#### Domain-level split-half reproducibility

To determine whether reproducibility extended beyond individual subscales to higher-order behavioral organization, split-half associations were additionally summarized at the domain level. Predictors were grouped into four behavioral domains (Sensory, Motor, Social, and Repetitive/RRB) and domain × outcome signed √ΔR² estimates were calculated independently in the discovery and replication subsets. This produced 36 domain × outcome cells (4 domains × 9 outcomes). Domain-level reproducibility was quantified using *Pearson* correlation between discovery and replication signed √ΔR² estimates across these 36 cells. Discovery-and replication-half domain × outcome matrices were also examined separately to assess preservation of the higher-order phenotypic architecture.

#### Anxiety-adjustment sensitivity analysis

To assess the contribution of shared anxiety-related variance to the observed architecture, the primary models were re-estimated after including CBCL Anxious/Depressed as an additional covariate. Because Anxious/Depressed served as a covariate in these models, it was removed from the corresponding outcome set. For each eligible predictor-outcome combination, a baseline model and an anxiety-adjusted model were estimated. FDR correction was performed independently for the baseline and anxiety-adjusted analyses, and hubness was recalculated using the corresponding model-specific FDR-adjusted associations. Changes in signed √ΔR² were used to quantify attenuation of individual associations after anxiety adjustment. Persistence of the broader predictor hierarchy was evaluated by comparing baseline and anxiety-adjusted hub ranks using Spearman rank correlation [61]. Baseline and adjusted hubness were calculated over matched CBCL outcome sets so that changes in rank reflected adjustment rather than differences in outcome composition.

#### Age-stratified phenotypic architecture

To examine developmental differences in relative phenotypic organization, the SPARK architecture was estimated separately within four age groups: 4-8, 9-12, 13-17, and ≥18 years. Signed √ΔR², FDR correction, and hubness were calculated independently within each age group using the same predictor and outcome definitions as in the primary architecture. Developmental stability was quantified by Spearman correlation between the 10-predictor hub-rank vectors for each pair of age groups. These correlations measure preservation of the relative ordering of behavioral predictors rather than longitudinal within-person change.

#### Equal-sample-size sensitivity analysis

Because available sample size differed across developmental groups, an equal-*N* sensitivity analysis tested whether differences in statistical power could account for age-related rank instability. Age groups were repeatedly subsampled without replacement to a common sample size (*n* = 17,866), and the age-specific analysis was repeated across 20 iterations. Pairwise hub-rank correlations were recalculated in each iteration, and the distribution of Spearman correlations was summarized for developmental comparisons.

#### Independent replication in SSC

The primary architectural analyses were repeated independently in SSC using corresponding DCDQ, RBS-R, SCQ, and CBCL measures.

Signed √ΔR² effects were estimated using the same general modeling framework, followed by FDR correction and calculation of predictor-level Hubness Indices. External replication was assessed at two complementary levels. First, preservation of the higher-order predictor hierarchy was quantified using Spearman correlation between SPARK and SSC hub ranks. Second, cell-level replication was evaluated across the 60 predictor × CBCL associations available in both cohorts using Pearson correlation between SPARK and SSC signed √ΔR² estimates.

SSC hubness and signed √ΔR² matrices were additionally examined directly to characterize the replicated architecture within the independent cohort.

#### Developmental stability in SSC

To determine whether age-related changes in hub organization generalized across cohorts, SSC hubness was evaluated separately within the available 4–8, 9–12, and 13–17-year age groups. Pairwise Spearman correlations between age-specific predictor ranks were calculated using the same 10-predictor hierarchy. These analyses were interpreted as cross-sectional comparisons of age-stratified architecture rather than within-person developmental trajectories.

#### Developmental domain-psychopathology coupling

The age-stratified hub analysis quantifies changes in the **relative organization** of predictors. A separate set of analyses therefore examined developmental differences in the **absolute strength of association** between broad behavioral domains and psychopathology.

#### Domain composites

Behavioral measures were aggregated into Sensory-Repetitive (SR), Motor, and Social domain composites. Constituent measures were standardized before aggregation, and available standardized features within each domain were averaged to create participant-level composite scores. Participants with no available constituent measure for a given domain were treated as missing for that domain. The psychopathology outcome was a standardized CBCL composite constructed from the available constituent psychopathology scales. Analyses were performed across four developmental bands: 0-4, 4-8, 8-12, and 12-18 years. Motor estimates were not available for ages 0-4 because DCDQ measures were unavailable in this age range.

#### Domain-level signed √ΔR²

For each estimable domain × age-band combination, population-level coupling with psychopathology was quantified using the same signed

√ΔR² framework:

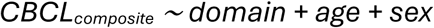

relative to:

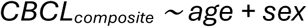

FDR correction was applied across the 11 estimable domain × age-band cells. Differences among domains were evaluated using weighted analysis of variance across the cell-level signed √ΔR² estimates. Domain-specific developmental trends and pairwise comparisons were used to characterize variation across age bands. Bayesian cell-level models provided complementary estimates of domain and developmental contrasts, summarized using posterior means, 95% credible intervals, and posterior probability of direction.

#### Clinical-threshold analysis

To examine whether the domain architecture extended to clinically elevated psychopathology, logistic regression modeled clinically elevated CBCL scores (T ≥ 63) as a function of standardized domain score, behavioral domain, developmental band, and their interactions, with age and sex included as covariates.

Domain × age-band effects were expressed as odds ratios with corresponding uncertainty intervals. This analysis provided a clinically interpretable complement to the continuous signed √ΔR² estimates.

#### Individual-level domain–psychopathology associations

To complement the population-level effect-size analysis, individual-level observations were stacked across available behavioral domains and modeled as a function of standardized domain score, domain, developmental band, and their interactions, with age and sex included as covariates.

Because individual participants could contribute observations for more than one behavioral domain, linear mixed-effects models with participant-specific random intercepts were used to account for within-person dependence. Domain-and age-specific slopes quantify the expected change in standardized psychopathology score associated with a 1-SD increase in the corresponding behavioral-domain composite. These analyses distinguish developmental variation in population-level architecture from variation in the individual-level association between behavioral burden and psychopathology.

#### Validation of domain aggregation

Domain-level architecture was compared with the corresponding predictor-level organization, and reproducibility of the aggregated domain structure was assessed using the independent SPARK split halves. These analyses tested whether the developmental domain representation retained the major organization evident at the finer subscale level.

#### Statistical analysis

All statistical tests were two-sided unless otherwise specified. Benjamini–Hochberg FDR correction was applied within each prespecified family of multiple tests, excluding unavailable/nonfinite tests from the denominator. *Pearson* correlation was used to quantify concordance of continuous effect-size estimates, whereas Spearman correlation was used for rank-based comparisons. Statistical analyses were conducted in Python using NumPy, pandas, SciPy, statsmodels, and scikit-learn.

Figures were generated using matplotlib and seaborn.

